# Bayesian modeling of mutual exclusivity in cancer mutations

**DOI:** 10.1101/2024.10.29.620937

**Authors:** Paweł Czyż, Niko Beerenwinkel

## Abstract

When cancer develops, gene mutations do not occur independently, prompting researchers to pose scientific hypotheses about their interactions. Synthetic lethal interactions, which result in mutually exclusive mutations, have received considerable attention as they may inform about the structure of aberrant biological pathways in cancer cells and suggest therapeutic targets. However, finding patterns of mutually exclusive genes is a challenging task due to small available sample sizes, sequencing noise, and confounders present in observational studies. Here, we leverage recent advancements in probabilistic programming to propose a fully Bayesian framework for modeling mutual exclusivity based on a family of constrained Bernoulli mixture models. By forming continuous model expansion within the iterative Bayesian workflow, we quantify the uncertainty resulting from small sample sizes and perform careful model criticism. Our analysis indicates that alterations in the EGFR and IDH1 genes may exhibit mutual exclusivity in glioblastoma multiforme tumors. We argue that Bayesian analysis offers a conceptual, systematic, and computationally feasible approach to model building, complementing the findings obtained from classical hypothesis testing approaches.

**Code:** https://github.com/cbg-ethz/jnotype

## 1 Introduction

Understanding gene interactions is fundamental in biology, particularly in cancer research: mutations at different loci often do not occur independently, but instead display patterns of cooccurrence and exclusivity (Canisius et al., 2016). The detection of mutually exclusive gene mutations is typically based on hypothesis testing (Vandin et al., 2011; Szczurek and Beerenwinkel, 2014; Constantinescu et al., 2015; Canisius et al., 2016; Liu et al., 2020; Shuaibi et al., 2024), where rejection of the null hypothesis is interpreted as evidence for mutual exclusivity.

Although hypothesis tests are efficient screening methods to employ on many possible gene sets under consideration, they compress the information about relative adequacy of two competing models into a single *p*-value and, subsequently, a dichotomous decision whether a pattern is deemed to be statistically significant or not.

Is it possible to extract more information from the data? Cumming (2013), Gelman and Shalizi (2013), Kruschke and Liddell (2018) and McElreath (2020) argue to shift the focus from rejection of a null hypothesis to building explicit models of the data generating process and estimating their parameters. In particular, this approach allows one to imagine what outcomes could be observed, if an experiment targeting genetic interactions was performed in a laboratory, and plan such experiments based on plausible observable outcomes, rather than relying solely on the statistical significance of the hypothesized pattern.

For finding patterns of mutual exclusivity, we consider the generative model of Szczurek and Beerenwinkel (2014). The authors propose estimating the parameters of two distinct generative processes, one generating patterns of mutual exclusivity and one background model, and selecting the model basing on Vuong’s closeness test (Vuong, 1989). Here, we propose to include both models in the iterative Bayesian workflow (Gelman et al., 2013; Blei, 2014; Betancourt, 2020; Gelman et al., 2020), which allows one to explicitly quantify uncertainty on the parameter estimates, and prioritize laboratory testing on hypothesized mutual exclusivity patterns basing on the plausible experimental outcomes, as predicted by the models.

However, predictions of any statistical method depend on whether the assumed model can adequately describe the data. We devise summary statistics to perform Bayesian model validation of the considered models. This approach allows us to find instances where neither of the models adequately describe the collected data. Following Draper (1995), Gelman and Shalizi (2013), and Gelman et al. (2020), we therefore seek suitable expansions of the existing models. Here, we show that both generative models (mutual exclusivity and background) are special instances of constrained Bernoulli mixture models (Xu, 2017). We leverage this observation to devise principled and scalable model expansions, in which Bayesian inference is performed with generic Hamiltonian Markov chain Monte Carlo samplers (Hoffman and Gelman, 2014; Betancourt, 2018).

### Contributions

Starting from the model of Szczurek and Beerenwinkel (2014), we analyze mutual exclusivity in cancer genotypes using a modern Bayesian workflow: We show how to explicitly quantify uncertainty in parameter estimates, provide a set of methods to assess when a model is adequate to describe observed data, and develop the framework for principled model expansion based on constrained Bernoulli mixture models. Constrained Bernoulli mixture models (Section 2.3) allow us to perform efficient inference in the presence of sequencing noise using generic Markov chain Monte Carlo methods, generate artificial observations, and remove sequencing noise from the collected data. Our analysis in Section 3 indicates that alterations in the EGFR and IDH1 genes may exhibit mutual exclusivity in a glioblastoma multiforme data set of Chang et al. (2013).

## 2 Methods

### 2.1 Mutual exclusivity as a tool guiding decision making

We consider a scientific hypothesis anticipating interactions within a small, prespecified set of genes. Such a gene set could be obtained through a large-scale screening effort using existing fast hypothesis tests on collected genomic data (Vandin et al., 2011; Szczurek and Beerenwinkel, 2014; Canisius et al., 2016; Liu et al., 2020), selected to represent a scientific hypothesis based on existing biological knowledge accessible via specialized databases (Kanehisa and Goto, 2000; Sondka et al., 2018), guided through other data modalities, such as cancer cell states obtained from single-cell transcriptomic data (Barkley et al., 2022), or selected solely through one’s intuition and experience.

The considered gene set has to be validated in the laboratory (Mani et al., 2008), as even the best statistical analysis based only observational data can be subject to confounding and data collection biases (Pearl, 2009; Schill et al., 2023). We therefore imagine a researcher planning to execute a series of perturbation experiments investigating the dependencies between the considered genes. How can one decide whether the particular gene set is worth investing time and laboratory resources, instead of using them to investigate alternative gene sets?

One possible method would be to use historical data to develop a model of the underlying data generating process. Such a model could, in principle, estimate expected effect sizes and potentially help determine the necessary sample size to gather sufficient evidence (Sankaran and Holmes, 2023). While this approach would require careful data modeling, it might be more time- and cost-effective than laboratory experiments. Additionally, it could offer insights into possible experiment outcomes, aiding researchers in prioritizing gene sets. Therefore, we are interested in creating a family of plausible generative processes to model the data and tentatively assess the strength of mutual exclusivity evidence in the available historical data.

### 2.2 Modeling noise

Assume that a gene set of size *G* has been proposed and consider the space of binary sequences 𝒴 = {0, 1}^*G*^. Let ***Y*** = (*Y*_1_, …, *Y*_*G*_) be a 𝒴-valued random vector representing a tumor genotype: *Y*_*g*_ = 1 when a mutation is present at locus *g* and *Y*_*g*_ = 0 otherwise. We are interested in modeling the distribution of ***Y***.

However, we cannot directly observe ***Y*** in historical data. Our primary assumption is that the errors arise independently, with the false positive rate *α* and false negative rate *β*, that is ℙ(*D*_*g*_ = 1 | *Y*_*g*_ = 0, *α*) = *α* and ℙ(*D*_*g*_ = 1 | *Y*_*g*_ = 1, *β*) = 1 − *β*, for each locus *g*. As the errors are assumed to arise independently, ***D*** | ***Y***, *α, β* ∼ Ber^*G*^ (*α* + (1 − *α* − *β*)***Y***), where Ber ^*G*^ (***p***) for ***p*** = (*p*_1_, …, *p*_*G*_) ∈ [0, 1]^*G*^ is the distribution on 𝒴 representing *G* independent Bernoulli trials with the corresponding probabilities *p*_1_, …, *p*_*G*_, respectively. We allow *p*_*g*_ = 0 and *p*_*g*_ = 1, representing Bernoulli trials with one-sided coins and use the notation *α* + *k****Y*** for the vector with components (*α* + *k****Y***)_*g*_ := *α* + *kY*_*g*_.

Apart from the noise model, we consider a parameter ***ψ*** ∈ Ψ, indexing possible probability distributions over the true (unobserved) genotype vector ***Y*** | ***ψ*** ∼ ℙ(***Y*** | ***ψ***) (see Fig. 1). There exist many plausible model classes for the distribution ℙ(***Y*** | ***ψ***), including various cancer progression models (Desper et al., 1999; Constantinescu et al., 2015; Schill et al., 2019). By combining the distribution ℙ(***Y*** | ***ψ***) with the noise model, we obtain a probability distribution over ***D***:

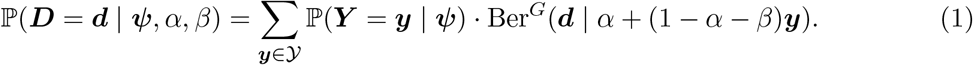

However, we do not have access to the distribution of ***D*** directly, but rather to a finite sample **𝒟** = ***D***_1_, …, ***D***_*N*_ ∈ 𝒴^*N*^, where *N* is the number of genotypes in the historical data set. We assume that the observed genotypes are conditionally independent and identically distributed given the parameters,

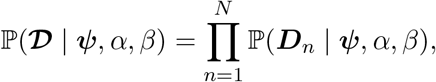

and we aim to infer model parameters ***ψ*** together with the noise rates *α* and *β*.

**Figure 1.**
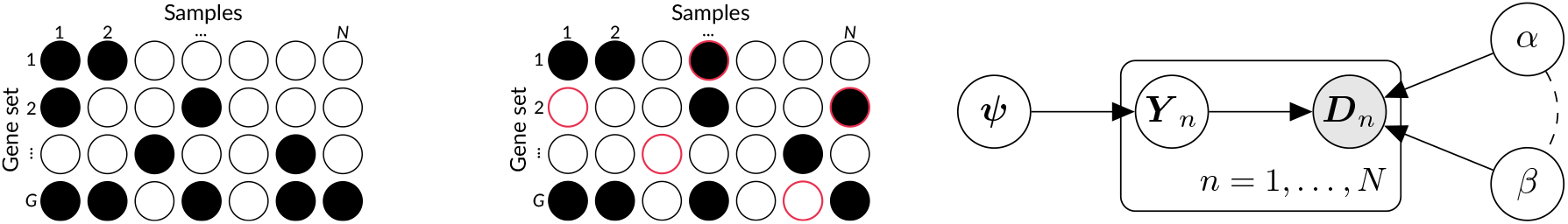
Tumor genotypes ***Y*** _*n*_ (left) are not directly observed. Instead, we observe the data subject to false positives and false negatives, ***D***_*n*_ (middle). All models considered in this work share the same graphical structure (right), linking unobserved tumor genotypes ***Y*** _*n*_ to the observed data ***D***_*n*_. Dashed lines signify possible dependencies between the false positive rate *α* and false negative rate *β*. Modelspecific parameter vector ***ψ*** defines the distribution over tumor genotypes.

We use Bayesian inference (Robert, 2007; Gelman and Shalizi, 2013; Gelman et al., 2013, 2020), which yields valid results even when considered models are non-identifiable, and allows us to incorporate domain knowledge by using the prior distribution ℙ(d***ψ*** d*α* d*β*) defined on the parameter space Ψ *×* (0, 1)^2^. By using generic Hamiltonian Markov chain Monte Carlo samplers (Hoffman and Gelman, 2014; Betancourt, 2018), we then obtain a sample from the posterior distribution ℙ(d***ψ*** d*α* d*β* | **𝒟**), explicitly quantifying how well different parameter settings agree with the observed data **𝒟** and our prior information. By comparing the predictions of the models under priors of varying strength, we can understand how robust our results are under different assumptions. We discuss further the employed Bayesian workflow in Section 2.5.

### 2.3 Constrained Bernoulli mixture models

Given a model ℙ(***Y*** |***ψ***), one has to marginalize out the variable ***Y*** as in Eq. 1, which generally requires 2^*G*^ summations, limiting the computational feasibility of modeling noise in the data (see Appendix A.3 for the computational complexity analysis). However, Bernoulli mixture models (Allman et al., 2009; Xu, 2017; Gu and Dunson, 2023; Malsiner-Walli et al., 2024) can be used to avoid this expensive summation. Write ***ψ*** = (***θ, ν***), where ***θ*** = (***θ***_1_, …, ***θ***_*K*_) ∈ [0, 1]^*K×G*^ is a matrix representing mixture components and ***ν*** ∈ Δ^*K*−1^ = {***v*** ∈ (0, 1)^*K*^ | *v*_1_ + … + *v*_*K*_ = 1} represents the mixture weights. The probability distribution ℙ(***Y*** | ***ψ***) is then given by

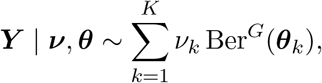

and can be calculated using *O*(*KG*) operations per unique genotype (see Appendix A.3). Computation of the observed likelihood function ℙ(***D*** | ***ψ***, *α, β*) does not increase the computational complexity:

**Lemma 1**. *Consider the probabilistic graphical model in Fig. 1 with* ***ψ*** = (***ν, θ***). *Let*

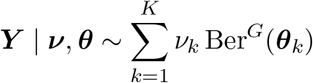

*be a binary vector distributed according to a Bernoulli mixture and* ***D*** | ***Y***, *α, β* ∼ Ber^*G*^((1 − *β*)***Y*** + *α*(1 − ***Y***)) *be a noisy version of it. Then*,

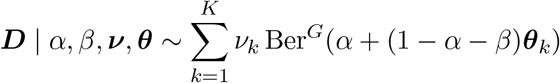

*is another Bernoulli mixture*.

This is not the most general form of the result: for a generalization and the proof, we refer to Appendix A.1. Alternatively, one can deduce it from the equivalent formula provided by Borgsmüller et al. (2020, Eq. 1). Hence, any (possibly constrained) Bernoulli mixture model for ***Y*** can be easily adapted to incorporate the noise. Conversely, having fitted a model to the observed data **𝒟**, we can sample noiseless genotypes ***Y*** and calculate the distribution of plausible future experimental outcomes. We further expand on these points in Section 2.4; however we first relate the framework of constrained Bernoulli mixtures to the two generative processes of Szczurek and Beerenwinkel (2014).

#### Background model

The background model for ***Y*** assumes that all loci mutate independently, i.e., ***Y*** | ***π*** = Ber^*G*^(***π***), where ***π*** ∈ (0, 1)^*G*^ is a parameter vector describing the mutation probabilities, ℙ(*Y*_*g*_ = 1 | ***π***) = *π*_*g*_. This is an example of a one-component Bernoulli mixture model, obtained by setting *K* = 1.

From Lemma 1 we see that the distribution of observed genotypes ***D*** depends only on the vector ***π****′* = *α* + (1 − *α* − *β*)***π***, with *π′*_*g*_ representing the probability of observing a mutation at locus *g*, i.e., ℙ(*D*_*g*_ = 1 | ***π***, *α, β*) = *π′*_*g*_. In particular, the parameters *α, β* and ***π*** are not jointly identifiable from the model.

#### Rigid mutual exclusivity model

As an alternative model, Szczurek and Beerenwinkel (2014) propose to estimate noise rates *α* and *β* together with a parameter vector ***ψ*** = (*γ, ξ*) ∈ (0, 1)^2^ as in Fig. 1, which parameterize the distribution of ***Y*** in the following manner. First, a latent subpopulation indicator is drawn from the categorical distribution with classes {1, 2, …, *G*+ 1} and probabilities *Z* | ***ψ*** ∼ Cat(*γ/G*,…, *γ/G*, 1 −*γ*). Then, if *Z* = *G* + 1, no mutation occurs: (***Y*** | ***ψ***, *Z* = 0) = **0**. However, if *Z* ∈ {1, …, *G*} (which occurs with *coverage probability γ*), then the *Z*th loci is exclusively mutated, meaning that *Y*_*Z*_ = 1 and, for all *g* ≠ *Z*, each locus mutates independently with chance *Y*_*g*_ | *Z*, ***ψ*** ∼ Ber(*ξ*). Hence, when the *impurity parameter ξ* is close to zero, we have strong mutual exclusivity, where only a single gene can be mutated. Larger values of *ξ* allow for some degree of co-occurrence. Similarly, small values of the coverage parameter *γ* signify that most samples ***Y*** _*n*_ are not exclusively mutated and the observed alterations in ***D***_*n*_ are more likely to occur due to the false positive rate *α*.

By marginalizing out the variable *Z*, we see that this is a constrained Bernoulli mixture model with *K* = *G* + 1 components:

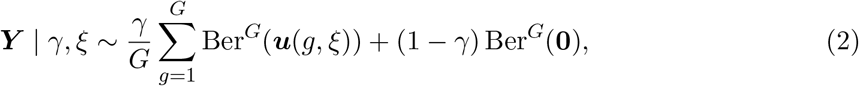

where *u*_*h*_(*g, ξ*) = 𝟙 [*g* ≠ *h*]+*ξ* 𝟙 [*g* ≠ *h*] is the probability of gene *h* mutating when gene *g* is chosen as exclusively mutated (i.e., *Z* = *g*). The indicator function 𝟙 [*φ*] evaluates to 1 if formula *φ* is true and to 0 otherwise.

Is this model identifiable? Lemma 1 shows that when *α* + *β* = 1, the probability ℙ(***D*** | *α, β, γ, ξ*) does not depend on *γ* and *ξ* anymore. This issue can be resolved by using a prior allowing only parameters such that *α* + *β* < 1. In fact, Szczurek and Beerenwinkel (2014) prove that for *G* ≥ 3 this model is generically locally identifiable (see Appendix A.4) and propose to set constraints on the noise rates, such as *β* = 0 or *α* = *β* = 0. Using an informative Bayesian prior, which puts most of the probability mass around small values of *α* and *β*, may therefore be an alternative to weaken this assumption.

#### Flexible mutual exclusivity model

Note that in the model above, the probability ℙ(*D*_*g*_ = 1 | *α, β, γ, ξ*) is the same for all loci *g* and the model is not flexible enough to model gene sets with differing observed mutation rates. While Szczurek and Beerenwinkel (2014) use this fact to develop an efficient hypothesis testing scheme, they also mention that one may prefer to relax this constraint. We propose to extend the model to

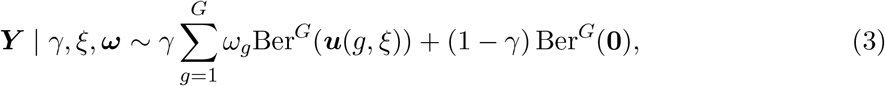

where we use the Dirichlet prior 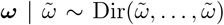 to control how balanced the patterns are expected to be. In particular, by setting a very strong prior (with 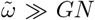), we recover the rigid model, in which *ω*_*g*_ ≈ 1*/G* for all *g*. Hence, this is an example of a continuous model expansion (Draper, 1995; Gelman and Shalizi, 2013), which is provided within the framework of constrained Bernoulli mixtures.

#### Joint model expansion

Finally, we build a model which simultaneously extends both the background and the mutual exclusivity models. Consider a vector ***π*** ∈ (0, 1)^*G*^ and a mixture

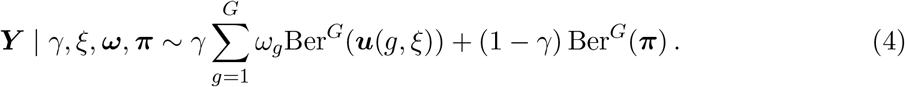

When *π*_*g*_ = 0 for all loci *g*, we obtain the flexible mutual exclusivity model (which then reduces to the rigid one for *ω*_*g*_ = 1*/G*). Instead, when *γ* = 0, we obtain the background model. Thus, this mixture model is a continuous model expansion of both previous models and the coverage parameter *γ* now explicitly controls how much mutual exclusivity is needed to explain the observed data.

### 2.4 The predictive distribution

As discussed above, Markov chain Monte Carlo methods allow us to efficiently sample from the posterior of the model parameters ℙ(d***ψ*** d*α* d*β* | **𝒟**). We can, however, obtain more information about the system by studying the predictive distributions ℙ(***Y*** | **𝒟**) (and, analogously, ℙ(***D*** | **𝒟**)). Note that as ℙ(***Y*** | **𝒟**) = ∫ ℙ(***Y*** | ***ψ***) ℙ(d***ψ*** | **𝒟**), we can sample a genotype vector ***Y*** * from the predictive distribution ℙ(***Y*** | **𝒟**) by reusing a sample ***ψ***\* ∼ ℙ(d***ψ*** | **𝒟**) from the posterior, and then sampling ***Y*** * from the mixture distribution ℙ(***Y*** | ***ψ***\*). Similarly, given an observed noisy observation ***D***, we can impute plausible values for the corresponding noiseless genotype vector ***Y*** by sampling from the conditional distribution ℙ(***Y*** | ***D*, 𝒟**) = ∫ ℙ(***Y*** | ***D, ψ***, *α, β*) (d***ψ*** d*α* d*β* | **𝒟**) in an analogous manner.

Moreover, Bernoulli mixture models have a structure allowing us to analytically extract meaningful summary statistics of the predictive distribution ℙ(***Y*** | ***ψ***) for each parameter setting ***ψ***. For example, Shuaibi et al. (2024, Sec. 2.2) argue to measure mutual exclusivity between two loci *g* and *h* by the log-odds ratio (LOR)

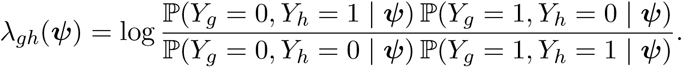

If, for parameter settings ***ψ***, the mutations arise independently, we have *λ*_*gh*_(***ψ***) = 0, while in the presence of mutual exclusivity we expect *λ*_*gh*_(***ψ***) *>* 0. While the probability ℙ(*Y*_*g*_ = 0, *Y*_*h*_ = 1 | ***ψ***), in principle, can be estimated by sampling sufficiently many artificial genomes ***Y*** * ∼ ℙ(***Y*** | ***ψ***), in Appendix A.2 we show how obtain this quantity analytically at *O*(*K*) cost.

Hence, as we have access to the samples from the posterior distribution ***ψ***\* ∼ ℙ(d***ψ*** | **𝒟**), we can compute the sample from the predictive distribution over the log-odds ratio, by analytically evaluating *λ*_*gh*_(***ψ***\*). This predictive distribution explicitly quantifies uncertainty: if we see that this predictive distribution is near 0, we can expect that mutual exclusivity may be rather weakly supported by historical data and prefer to prioritize other experiments.

This is not the only possible measure of mutual exclusivity. As another example, consider the difference of conditional expectations (DCE), Δ_*g*|*h*_(***ψ***) = 𝔼[*Y*_*g*_ | *Y*_*h*_ = 1, ***ψ***] − 𝔼[*Y*_*g*_ | *Y*_*h*_ = 0, ***ψ***]. If *Y*_*h*_ was a variable assigned to a controlled intervention (as in a perturbation experiment), rather than merely an observational quantity, DCE would correspond to the average treatment effect of *Y*_*h*_ onto *Y*_*g*_. However, this quantity does not have explicit causal interpretation in our observational framework. Instead, Δ_*g*|*h*_ quantifies the change in the probability of mutation at locus *g* between subpopulations defined by the mutation status at locus *h*.

Other measures of co-occurrence and mutual exclusivity are also possible: we discuss them in Appendix A.2.1. More generally, if *f* : Ψ → ℝ is any measurable function representing a quantity of interest (Lundberg et al., 2021), we can construct its posterior predictive distribution using the collected samples ***ψ***\* ∼ ℙ(d***ψ*** | **𝒟**) and evaluating *f* (***ψ***\*) (Gelman et al., 2013, Ch. 5).

### 2.5 The Bayesian workflow

Following Gelman and Shalizi (2013) and Gelman et al. (2020), we proceed within the Bayesian workflow, where multiple models are used to perform the inference from the same data set. We then understand the possible data generating process by comparing predictions of different models and validating them against the collected data using graphical posterior predictive checks (Gelman et al., 2013, Ch. 6).

In particular, we do not attempt to perform a formal model selection procedure, e.g., by the means of Bayes factors (Robert, 2007, Ch. 5). We acknowledge the possibility that there may be multiple models fitting the available data reasonably or that neither of the considered models may offer a reasonable fit. Bayesian workflow performed in this manner, serves therefore a role outlined in Section 2.1: at the cost of additional effort spent on explicit modeling, we can learn more from the available data than by employing a hypothesis test, which selects one of two prescribed models.

To perform graphical posterior predictive checking, we choose a data statistic ***T*** : 𝒴^*N*^ → ℝ^*k*^, which maps the observed data set **𝒟** to a vector in some Euclidean space ℝ^*k*^, where *k* is referred to as the dimension of the statistic. We use the predictive distribution ℙ(***D*** | **𝒟**) (see Section 2.4) to generate multiple artificial data sets 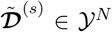 of the original size *N*. By visualizing the distribution of summary statistic 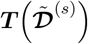 against ***T*** (**𝒟**), we can assess in which aspects the model underfits the data.

As summary statistics ***T*** for our mutual exclusivity model (Eq. X), we propose the following:

1. A *G*-dimensional statistic measuring observed mutation frequency at each locus, 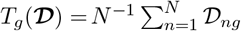
2. A (*G* + 1)-dimensional statistic measuring the fraction of samples with a given number of mutations: For *m* ∈ {0, 1, …, *G*}, define 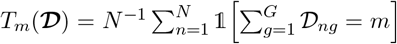. In the terminology of Canisius et al. (2016), 1 − *T*_0_ is named the coverage statistic, *T*_1_ is named exclusivity statistic, and the sum *T*_2_ + *T*_3_ + … + *T*_*G*_ is called impurity.
3. A *G*(*G* − 1)*/*2-dimensional statistic describing the observed Matthews correlation coefficient (MCC) between ***D***_*g*_ and ***D***_*h*_ for each (unordered) pair of distinct loci *g* ≠ *h*. As MCC is formally not defined when either data column is constant, in these cases we replace it by 0.

## 3 Results

In this section, we demonstrate how to employ the proposed approach to model several candidate mutual exclusivity patterns. We analyze the glioblastoma multiforme dataset from Chang et al. (2013), which consists of *N* = 236 samples. Szczurek and Beerenwinkel (2014) screened this dataset for statistically significant gene sets of size *G* = 4. They identified four gene sets: *A* (consisting of EGFR, GCSAML, IDH1, OTC), *B* (ABCC9, PIK3CA, RPL5, TRAT1), *C* (PIK3C2G, PIK3CA, RPL5, TRAT1), and *D* (NF1, PIK3C2G, PIK3R1, TRAT1). We visualize these gene sets in Fig. 2.

**Figure 2.**
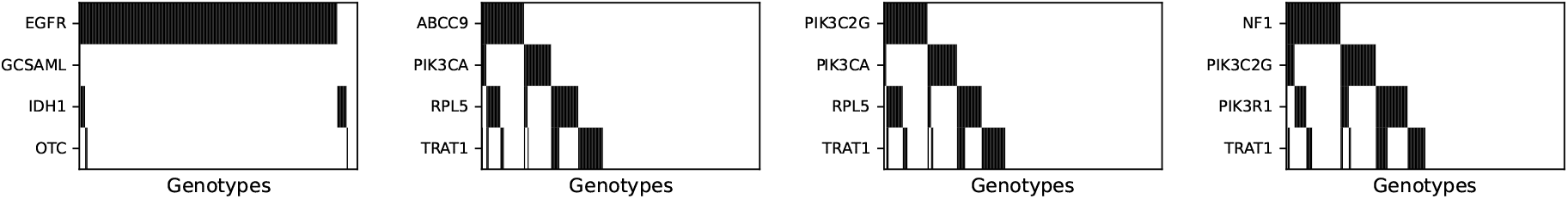
Four gene sets representing alterations in glioblastoma multiforme.

Gene sets *B, C*, and *D* were marked as statistically significant according to their mutual exclusivity test, with noise rates constrained to zero (*α* = *β* = 0). Gene set *A* was not selected as statistically significant according to the mutual exclusivity test, but was marked as statistically significant by the adjusted permutation test (Vandin et al., 2011). However, when Constantinescu et al. (2015) performed the mutual exclusivity test on multiple gene set sizes, and adjusted the results for multiple comparisons, no gene set of size four was marked as statistically significant.

These findings seem to support the point of view of Gelman and Stern (2006) and Imbens (2021) that *p*-values should be treated with appropriate caution. In particular, the statistical significance claims can strongly depend on the type of the hypothesis test used, applied multiple testing correction, and possibly the enforced constraints on the selected parameters. Below we show how to use the Bayesian workflow to re-analyze these gene sets and understand which of them could be the most promising to test experimentally.

### Model implementation

To perform Bayesian analysis, we implemented the joint model expansion (Eq. 4) in NumPyro (Phan et al., 2019), which contains the vector-valued parameters ***ν, ω*** ∈ Δ^*G*−1^ and ***π*** ∈ (0, 1)^*G*^, as well as scalar parameters *α, β, γ, ξ* ∈ (0, 1). We obtained approximations to the background model and mutual exclusivity models given by Eq. 2 and Eq. 3 by setting strong priors which constrained selected parameters. As all parameters are continuous, and JAX (Bradbury et al., 2018) provides automatic differentiation capabilities, we could use Hamiltonian Markov chain Monte Carlo algorithms, which employ the gradient of the loglikelihood to devise proposals with low autocorrelation and maintain high acceptance probability. Namely, we used the NUTS sampler (Hoffman and Gelman, 2014) and approximated the posterior distribution using four Markov chains. Within each chain, we used the warm-up phase to 8,000 samples (which were used to tune the sampling parameters), and then collected additional 8,000 samples. To spot possible convergence issues, we monitored the number of divergences which occurred during sampling (Betancourt, 2018), as well as calculated the potential scale reduction factor 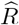 and the effective sample size (Gelman et al., 2013, Ch. 11).

### 3.1 Analysis of gene set *A*

We started by fitting the background model. We did not observe any convergence issues, with no sampler divergences, all 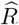 values below 1.01, and minimum effective sample size exceeding 27,000. However, by looking at the posterior predictive distribution of the MCC statistic, we noted that the strong negative correlation observed between mutations EGFR and IDH1 cannot be attributed to noise in this model (Appendix B.1).

Then, we fitted the rigid mutual exclusivity model, targeting balanced patterns (Eq. 2). Similarly, we did not see the convergence issues during sampling. However, the model severely underfits both MCC and the mutation frequencies (Appendix B.1): as the mutations are symmetric in this model, both mutation frequency and MCC between any pair of genes has to be the same.

Therefore, we fitted the more flexible version of the mutual exclusivity model (Eq. 3), allowing ***ω*** to vary. We used a weakly informative prior ***ω*** ∼ Dir(2, 2, …, 2) and, as in previous cases, did not observe convergence issues. The posterior on the coverage concentrates in the region close to 1 (Fig. 3a) and the model learns mutation frequencies precisely (Fig. 3b). Moreover, by investigating the predictive distribution of the LOR (*λ*; Fig. 3c) and DCE (Δ; 3d), we found that exclusivity between genes EGFR and IDH1 is supported by this model: the log-odds ratio attains values in range 4–6, agreeing with mutual exclusivity between these two mutations, while DCE attains negative values and does not include zero. We note, however, that the background model also provides a reasonable fit to the data, excluding presence of any interactions.

**Figure 3.**
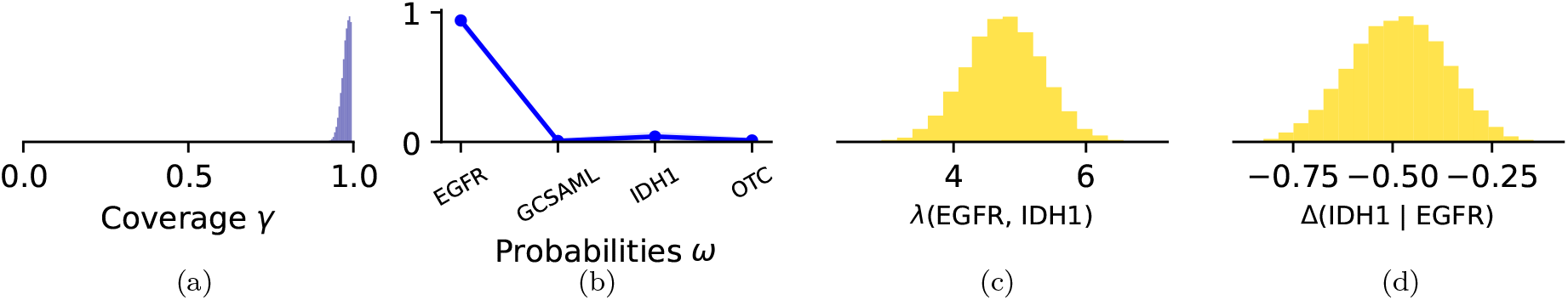
Flexible mutual exclusivity model fitted to gene set *A*.

Finally, we attempted to fit the full model (Eq. 4) varying the strength of the prior on the *γ* parameter: we used the spike-and-slab priors build using a mixture of two truncated normal distributions, where the spike component attributes large probability mass to the *γ* ≈ 0 region, preferring the background model. Namely, we used *γ* ∼ *p* 𝒯 𝒩 (0, 0.01; *ϵ*, 1 − *ϵ*) + (1 − *p*) 𝒯 𝒩 (0, 1; *ϵ*, 1 − *ϵ*) (where the truncation parameter *ϵ* = 0.005 is used to avoid potential numerical issues with sampling *γ* too close to 0 or 1). We varied the mass of the spike component within *p* ∈ {0.02, 0.25, 0.5}, where increasing *p* results in stronger preference of the background model over the mutual exclusivity model. Intuitively, if the evidence for mutual exclusivity is strong in the data, we should observe the posterior on the parameter *γ* being able to escape the penalization introduced by the spike component. Hence, by varying *p* and observing how the posterior on *γ* changes, we can understand how strong the signal in the data is. However, in this case we noticed convergence issues, potentially caused by multimodality of the posterior. We suspect that the Markov chain Monte Carlo sampler has trouble switching between modes corresponding to the background model and the flexible mutual exclusivity model.

### 3.2 Analysis of the gene sets *B, C*, and *D*

We fitted all the models to gene sets *B, C*, and *D* as described above. We did not spot any convergence issues in sampling.

In Fig. 4 we present the posterior predictive distributions of the flexible mutual exclusivity model fitted to gene set *B*. We see that the coverage attains values between 0 and 0.5, limiting the precision to which the proportion vector ***ω*** can be determined. However, we did not see strong evidence for mutual exclusivity: LOR *λ* attains mostly negative values (corresponding to co-occurrence), while DCE is positive. This behavior can be explained by a relatively large value of impurity parameter *ξ*. Hence, we note that the mutual exclusivity model may, in fact, support co-occurrence, and explicit investigation of the predictive distribution is crucial when planning future experiments targeting potential exclusivity.

**Figure 4.**
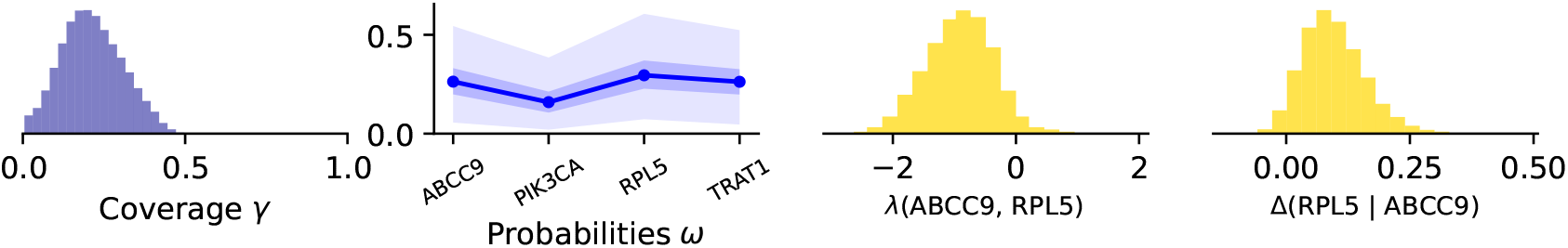
Flexible mutual exclusivity model fitted to gene set B.

In this case, we also compare this result with the predictions of the joint expansion model, with three different priors on the spike component (Fig. 5). We see that the model prefers to explain the data using the independent component, placing large probability mass around *γ* = 0. Subsequently, the log-odds ratio and conditional expectation difference is concentrated near zero. However, this model also suggests that co-occurrence is more plausible than mutual exclusivity (Appendix B.2). We draw similar conclusions for gene sets *C* and *D*, providing the detailed analysis in Appendices B.3 and B.4.

**Figure 5.**
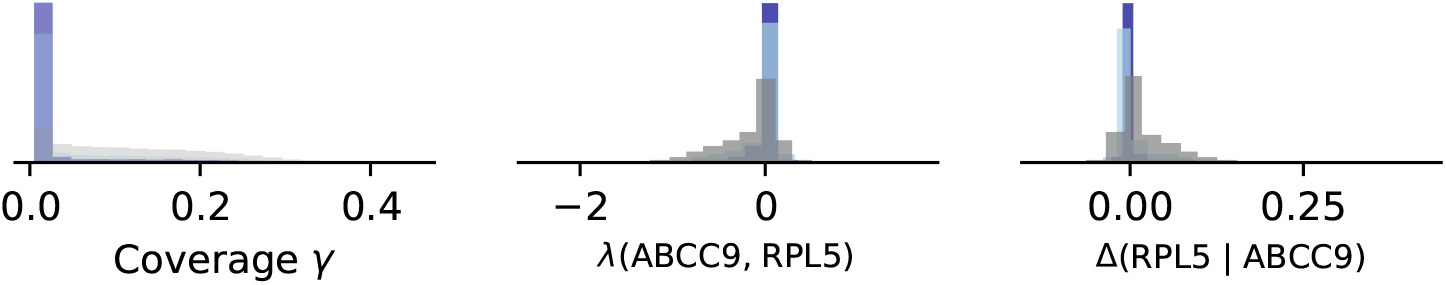
Joint model extensions with different priors on the coverage parameter *γ* fitted to the gene set

### 3.3 Reconciliation

By performing Bayesian analysis we noted that mutual exclusivity between alterations in EGFR and IDH1 may be plausible. On the other hand, the data can be reasonably well explained by the background model, which has vastly different predictions in this aspect. However, as we observed possible weak positive correlations, rather than exclusivity, in gene sets *B, C*, and *D*, we would prioritize executing the experiments involving gene set *A* over the alternatives.

We also noted unexpected possible positive correlations in gene sets *B, C* and *D* can be stronger than anticipated by the background model. This finding, can be due to random chance, but also due to possible hidden confounding by genetic alterations not included in the analysis of these particular gene sets or varying patient characteristic.

## 4 Discussion

In this manuscript, we argue for using Bayesian workflow to understand postulated mutual exclusivity patterns. Explicit Bayesian modeling allows us to quantify uncertainty on parameter estimates and draw insights from the predictive distributions of quantities of interest directly related to mutual exclusivity. We propose a flexible family of models, which are yet interpretable and convenient to fit to the data using modern Markov chain Monte Carlo methods. Finally, we discuss how to validate adequacy of the fitted models by comparing summary statistics against the observed data. While the proposed methodology is not free of challenges, which we further discuss below, Bayesian modeling can serve as an efficient intermediate step between exploratory hypothesis tests and extensive perturbation experiments, enabling the assessment of mutual exclusivity patterns and the prioritization of gene sets for further investigation.

### 4.1 Limitations and future work

#### The analysis is (and always will be) subjective

Bayesian workflow has many moving parts: we consider multiple models, each of them can be given different priors, and we look at different summary statistics. Moreover, there is no formally defined outcome of the analysis, which could be used to deterministically decide whether a particular gene set should be treated as significant or not. We do not feel particularly comfortable with so many subjective choices and see the risk that one could draw any conclusion from our analysis, by choosing a single model and using a strong prior. However, as discussed in Section 3, *p*-values are not objective either: they depend on the hypothesis test employed, chosen multiple testing correction, as well as the possible hypotheses which (even in principle) could have been executed (Gelman and Loken, 2019). In general, we cannot make biological discoveries from observational data alone: any insights obtained should be carefully validated by perturbation experiments. Performing the advocated analysis is supposed to help the researcher to navigate through the possible data generating processes and understand the strength of existing evidence for mutual exclusivity before executing a proper perturbation experiment.

#### Employing other generative models

While constrained Bernoulli mixture models are scalable and naturally model noise due to Lemma 1, there are many alternative choices for the model ℙ(***Y*** | ***ψ***) and when the gene set size *G* is sufficiently small, explicit summation in Eq. 1 can be scalable enough (cf. Appendix A.3). Therefore, one could incorporate more established models of binary genotypes into the Bayesian workflow, for example the mutagenetic trees framework (Desper et al., 1999), an Ising model (Xue et al., 2012), or various cancer progression models (Constantinescu et al., 2015; Schill et al., 2019; Greenbury et al., 2019). Additionally, Shuaibi et al. (2024) recently introduced a generative model of cancer mutations which extends beyond binary genotypes to the number of mutations occurring in a particular gene. Building a version of the Bayesian workflow for the count-based mutual exclusivity models may therefore be an interesting avenue for future research.

#### A large number of interrelated gene sets

In this work, we focus on Bayesian analysis of a single gene set at a time. However, the gene sets are selected from the historical data and when analyzing multiple gene sets, the data is reused multiple times (as a single gene may be investigated in multiple gene sets), which poses a methodological issue for our framework. Namely, when different gene sets are considered, we do not have a single generative process for the observed data, but rather we perform the analyses treating gene sets as fully independent.

A more principled approach could be based on building larger models, modeling simultaneously a larger number of genes (cf. Schill et al. (2019)). However, such models may be subject to model misspecification (Tarpey et al., 2008; Gelman and Shalizi, 2013; Watson and Holmes, 2016) or involve subtle non-identifiability and estimation issues (Gotovos et al., 2021; Schill et al., 2023). Larger models can also potentially help with adjusting the mutation data for patient-specific characteristics. For example, Bayer et al. (2023) incorporate clinical and demographic information into the model, to better untangle the existing patterns. The framework of constrained Bernoulli mixture models is especially convenient for incorporating such effects, becoming a dependent mixture model (see Wade et al. (2023) for a review).

#### Incorporating existing domain knowledge

Although historical data can guide experiments, in practice, a researcher will plan experiments using information from many different resources, such as existing databases containing known biological pathways (Kanehisa and Goto, 2000; Hanahan and Weinberg, 2011), interactions found in Cancer DepMap experiments (Tsherniak et al., 2017), or look for therapeutic potential (Waarts et al., 2022). Although in this manuscript we do not discuss techniques using information from data sets of different modalities, in principle, Bayesian approach is suited to borrowing information across experiments, e.g., by employing power priors (Ibrahim et al., 2001, Sec. 1.7). Although this manuscript does not explore techniques that integrate information from datasets of different modalities, Bayesian approaches are well-suited for borrowing information across experiments (Ibrahim et al. (2001, Sec. 1.7) and Frazier and Nott (2024)).

In conclusion, Bayesian workflow, grounded in principled model building and validation, enables the extraction of additional insights from available data. By utilizing a family of models with interpretable parameters and constructing predictive distributions, we can consider plausible outcomes for future experiments, representing a step towards the principled discovery of genetic interactions.

## Reproducibility statement

The code is available as a subpackage of the Jnotype framework (https://github.com/cbg-ethz/jnotype). We ensured reproducibility of our results by implementing them as Snakemake workflows (Mölder et al., 2021).

## Acknowledgments

Section 3 presents results based upon data generated by the TCGA Research Network: https://www.cancer.gov/tcga. PC is funded through the ETH AI Center fellowship program.

## Appendix

## A Technical results

### A.1 Proof of Lemma 1

Consider the following generalization of Lemma 1:

**Lemma 2**. *Consider a probabilistic graphical model in Fig. 1 with* ***ψ*** = (*F*, ***θ***), *where F is a probability measure on a standard Borel space* 𝒵 *and* ***θ*** : 𝒵 → [0, 1]^*G*^ *be a measurable function. Let*

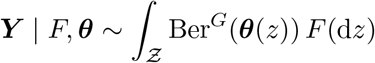

*be a binary vector distributed according to a Bernoulli mixture and*

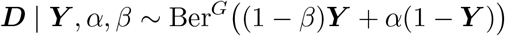

*be a noisy version of it. Then*,

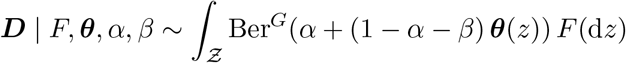

*is another Bernoulli mixture*.

For a finite space 𝒵 = {1, 2, …, *K*} and the discrete distribution *F* = *ν*_1_*δ*_1_ + … + *ν*_*K*_*δ*_*K*_, we obtain Lemma 1 as a special case. Below we prove the general version.

*Proof*. Let *Z* | *F* ∼ *F* represent the mixture component, so that ***Y*** | *Z*, ***θ*** ∼ Ber^*G*^(***θ***(*Z*)). We have

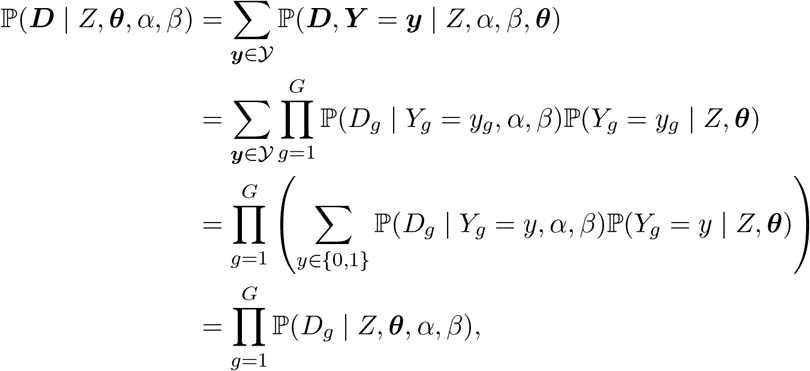

where each distribution ℙ(*D*_*g*_ | *Z*, ***θ***, *α, β*) is supported on the set {0, 1}, so that it has to be a Bernoulli distribution. Using the law of total expectation, we obtain

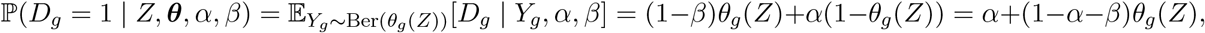

which gives

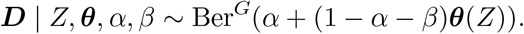

By integrating out the variable *Z*, we obtain the required mixture model.

To provide more intuition how the proof above mitigates the summation of 2^*G*^ terms corresponding to different elements of 𝒴 = {0, 1}^*G*^, consider the following Bayesian network:

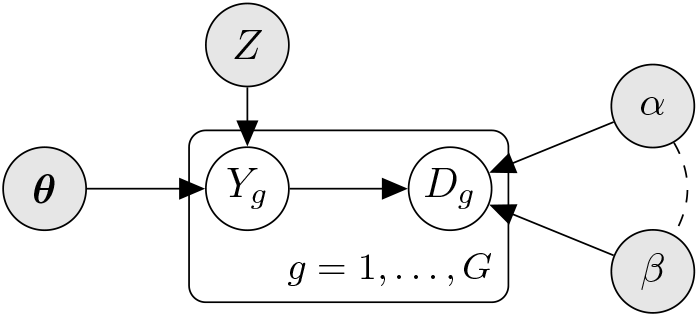

As we condition on the filled nodes, the tuples (*Y*_*g*_, *D*_*g*_) for distinct *g* become independent of each other, which is easy to see from the *d*-separation criterion (Pearl, 2009, Sec. 1.2.3) and a simple inductive argument. As the binary variables *D*_*g*_ are independent of each other, we can conclude that they jointly have to come from a suitable Ber^*G*^ distribution, where the probability vector can be obtained by marginalizing out *Y*_*g*_ variables in parallel, requiring *O*(*G*) (rather than *O*(2^*G*^)) summations.

This argument also shows that Lemma 2 is not the strongest version possible: one can introduce additional dependencies between the filled nodes, making the noise rates *α* and *β* dependent on the mixture component *Z*. This, however, would not correspond to the sequencing noise model studied in this work.

**Equivalent formula** Borgsmüller et al. (2020, Eq. 1) use a formula

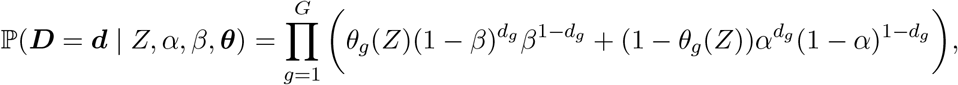

which is equivalent to Ber^*G*^(***d*** | *Z, α, β*, ***θ***). To see this, note that for every *p* ∈ [0, 1] and *d* ∈ {0, 1} one has

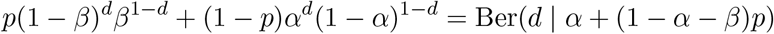

as for *d* = 1 both sides reduce to *p*(1 − *β*) + (1 − *p*)*α* = *α* + (1 − *α* − *β*)*p* and for *d* = 0 both sides reduce to *pβ* + (1 − *p*)(1 − *α*) = 1 (− *α* + (1 − *α* − *β*)*p*).

### A.2 Arbitrary conditioning in finite Bernoulli mixture models

In this section we expand on the point from Section 2.4, formally proving that finite Bernoulli mixture models have an arbitrary conditioning property. Hence, they belong to the family known as orderless autoregressive models (Uria et al., 2016) and were employed as baselines for such problems (ibid). This property follows from interpreting a finite Bernoulli mixture model as a member of a more general family of mixture models studied by Czyz? et al. (2023), but for completeness we present the proof here.

Let *I* and *J* be two disjoint sets of loci and ***y*** ∈ 𝒴 be any vector. We write ***Y*** _*I*_ for the random vector (*Y*_*i*_)_*i*∈*I*_. We want to efficiently calculate the conditional probability

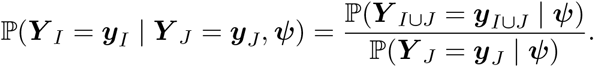

Note that it therefore suffices to calculate the marginal probabilities ℙ(***Y*** _*A*_ = ***y***_*A*_ | ***ψ***) for two sets of loci, *A* = *I* ∪ *J* and *A* = *J*.

For a single component this marginalization is easy due to conditional independence:

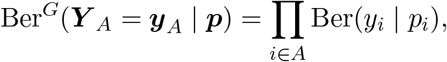

and can be accomplished in *O*(|*A*|) time, which never exceeds *O*(*G*). In a finite Bernoulli mixture model (with *K* components), we have

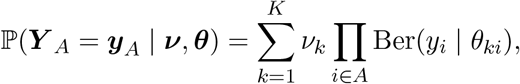

which can be calculated in *O*(*K* · |*A*|) time, which never exceeds *O*(*KG*) time.

Similarly as in Appendix A.3, it is beneficial for numerical stability to work with log-probabilities, rather than probabilities.

#### A.2.1 Measures of mutual exclusivity

The construction above allows us to obtain the value ℙ(*Y*_*g*_ = *y*_*g*_, *Y*_*h*_ = *y*_*h*_ | ***ψ***) and, similarly as in Section 2.4, to construct the predictive distribution by using samples from the posterior. For example, consider the difference of conditional expectations

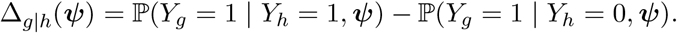

(We avoid the term “average treatment effect” as *Y*_*h*_ is not a controlled and randomly assigned treatment, but merely an observational quantity). This is an interesting estimand, although this is not the only choice possible and may suffer from the fact that the mutation probabilities, and their difference, can be rather small. As an alternative measure of exclusivity, we can consider the logarithmic probability ratio

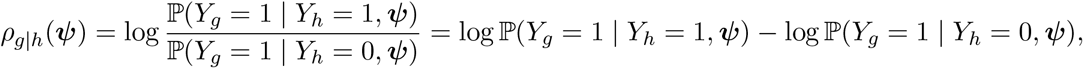

which may be easier to use where the differences in probabilities are very small. However, using the logarithm may amplify any noise.

Shuaibi et al. (2024) use the log-odds ratio,

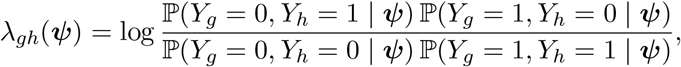

which is symmetric in indices *g* and *h* and which positive values may be indicative of mutual exclusivity.

Yet another measure of dependency between genes, although not sensitive enough to distinguish between mutual exclusivity and co-occurence, is the mutual information (Polyanskiy and Wu, 2022, Ch. 4), defined as the Kullback–Leibler divergence KL(ℙ(*Y*_*g*_, *Y*_*h*_) ∥ ℙ(*Y*_*g*_)ℙ(*Y*_*h*_)), which predictive distribution can be constructed as in Czyz? et al. (2023): for each sample from the posterior ***ψ*** we construct the mutual information KL(ℙ(*Y*_*g*_, *Y*_*h*_ | ***ψ***) ∥ ℙ(*Y*_*g*_ | ***ψ***)ℙ(*Y*_*h*_ | ***ψ***)).

### A.3 Efficient evaluation of the likelihood

We follow the usual convention and work with loglikelihood, rather than likelihood, which helps to prevent numerical errors coming from operations on small probabilities. Recall that the loglikelihood in the Ber^*G*^(***π****′*) model is given by

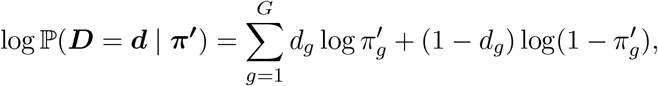

which requires *O*(*G*) operations. Due to Lemma 1, the likelihood in the Bernoulli mixture model can be evaluated as

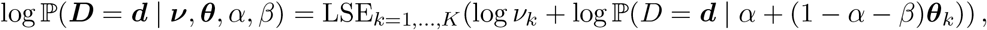

where LSE(*x*_1_, …, *x*_*K*_) = log(exp(*x*_1_) + … + exp(*x*_*K*_)) is the numerically stable logsumexp function. Typically, it is implemented as

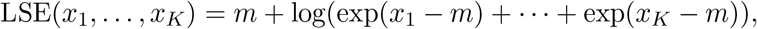

where subtracting *m* = max(*x*_1_, …, *x*_*K*_) prevents numerical overflow.

Note that the likelihood can therefore be calculated using *O*(*KG*) operations. As in the considered models we have *K* = *O*(*G*) components, the evaluation of the likelihood is polynomial in *G*. We envision that it is possible to add more components, parameterizing pairs of genes.

The loglikelihood evaluation over the whole data set takes, therefore, *O*(*NKG*) operations, which are however possible to parallelize over the *N* patients. However, for small gene sets, it is possible to additionally improve the computational performance. Let *U* be the number of unique genotypes in the observed sample. We have *U* ≤ min (*N*, 2^*G*^ ). We can calculate the loglikelihood for only *U* genotypes and construct the loglikelihood over the whole data set using a weighted sum of loglikelihoods corresponding to different unique genotypes. Hence, it is possible to reduce the computational complexity from *O*(*NKG*) to *O*(*UKG*).

Consider now a general model, in which a single call to the likelihood function ℙ(***Y*** | ***ψ***) requires *T* operations. When *α* = *β* = 0, with probability one it holds that ***D*** = ***Y***, and one can therefore calculate the likelihood on the whole data set in *O*(*UT*) operations. However, when the noise rates are non-zero, one has to explicitly marginalize ***Y*** as in Eq. 1. Hence, the overall complexity of the algorithm requires *O (T*2^*G*^) operations for calculating the likelihood on all the possible values of the noiseless genotype vector ***Y*** and an additional *O (GU*2^*G*^) factor for the calculation of the likelihood on the observed genotypes **𝒟**. Overall time complexity is therefore *O* (*GU* + *T*)2^*G*^), which is limiting for large gene sets, for which *N* ≪ 2^*G*^.

### A.4 Discussion of the identifiability of the mutual exclusivity model

Let Θ ⊆ ℝ^*k*^ which represents all model configurations ***θ*** = (***ψ***, *α, β*). We assume that Θ is measurable and has strictly positive Lebesgue measure. If the mapping 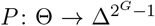 given by ***θ*** → ℙ(***D*** | ***θ***) is injective, one can recover model parameters having access to a sufficiently large data sample. This condition is known as strict identifiability (Allman et al., 2009).

However, strict identifiability of a Bernoulli mixture models is rarely possible (Gyllenberg et al., 1994; Allman et al., 2009; Najafi et al., 2020). Hence, Allman et al. (2009, p. 8) introduce the notion of of generic identifiability: a property is said to hold generically, if it holds for all elements in a subset Θ \ *V* (*f*_1_, …, *f*_*n*_), where

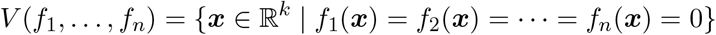

is a real algebraic variety given by the set of simultaneous zeros of polynomials *f*_1_, …, *f*_*n*_.

Note that an algebraic variety is a closed set, and hence is of Lebesgue measure zero. Therefore, if the parameters of the ground-truth data generating process are drawn from some distribution ℙ(***θ***) defined on Θ, which has density with respect to the Lebesgue measure, with probability one they have to land in a subset Θ \ *V* (*f*_1_, …, *f*_*n*_).

Allman et al. (2009) establish generic identifiablity for several Bernoulli mixture models. However, the question of generic identifiability for constrained Bernoulli mixture models remains an active area of research (Xu, 2017; Gu and Dunson, 2023; Liu and Culpepper, 2024). Note that every Bernoulli mixture model in Fig. 1 reduces to Ber^*G*^(*α, α*, …, *α*) for *α* + *β* = 1. Hence, the mutual exclusivity model from Section 2.3 cannot be strictly identifiable unless additional constraints are introduced (e.g., a natural constraint *α* + *β* < 1) or locally identifiable, which would mean that for every point there would exist a sufficiently small neighborhood on which the mapping *P* would be injective.

However, Szczurek and Beerenwinkel (2014, Supplement S1) prove that for *G* ≥ 3, the mapping 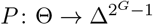 is generically immersive, meaning that its differential *P*′(***θ***) is injective for all points ***θ*** ∈ Θ\*V* (*f*_1_, …, *f*_*n*_). Hence, for every such point, there exists an open ball *B*_***θ***_ ⊂ Θ such that the restriction 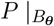 is injective. Therefore, the mutual exclusivity model is generically locally identifiable. However, it remains an open question whether the mapping is generically (globally) identifiable.

#### Consequences for Vuong’s closeness test applicability

As the mutual exclusivity model is not strictly identifiable, Assumptions A3 and A5 of Vuong (1989) may require suitable reformulation to ensure that Theorem 5.2 (ibid) holds. For example, note that for *α* + *β* = 1, the background and the mutual exclusivity model are not strictly non-nested, as the distribution Ber^*G*^(*p, p*, …, *p*) is included in both models. In this case, a different asymptotic distribution of the likelihood ratio statistic (than for strictly non-nested models) may arise (Wilson, 2015).

## B. Additional experimental information

In this section, we present information supplementing the analysis in Section 3.

### B.1 Analysis of the gene set *A*

The posterior predictive checks in Fig. 7 show that both background model and rigid mutual exclusivity model underfit the provided data. The flexible mutual exclusivity model (Eq. 3) better captures the distribution. As we see in Fig. 6, the flexible mutual exclusivity model suggests that genes IDH1 and EGFR may be mutually exclusive.

**Figure 6.**
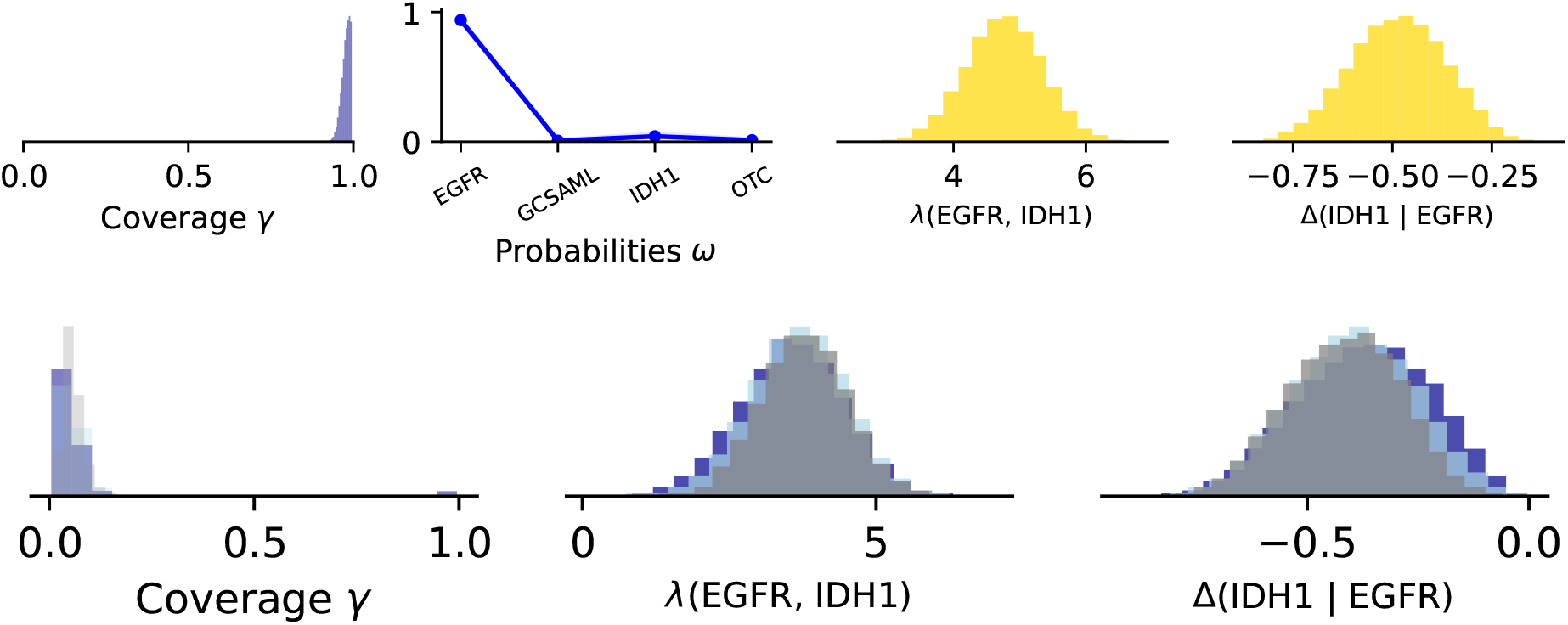
Predictions from different models applied to gene set *A*. Top row: flexible mutual exclusivity model (Eq. 3). Bottom row: full model (Eq. 4) with different priors on the coverage parameter *γ*.

**Figure 7.**
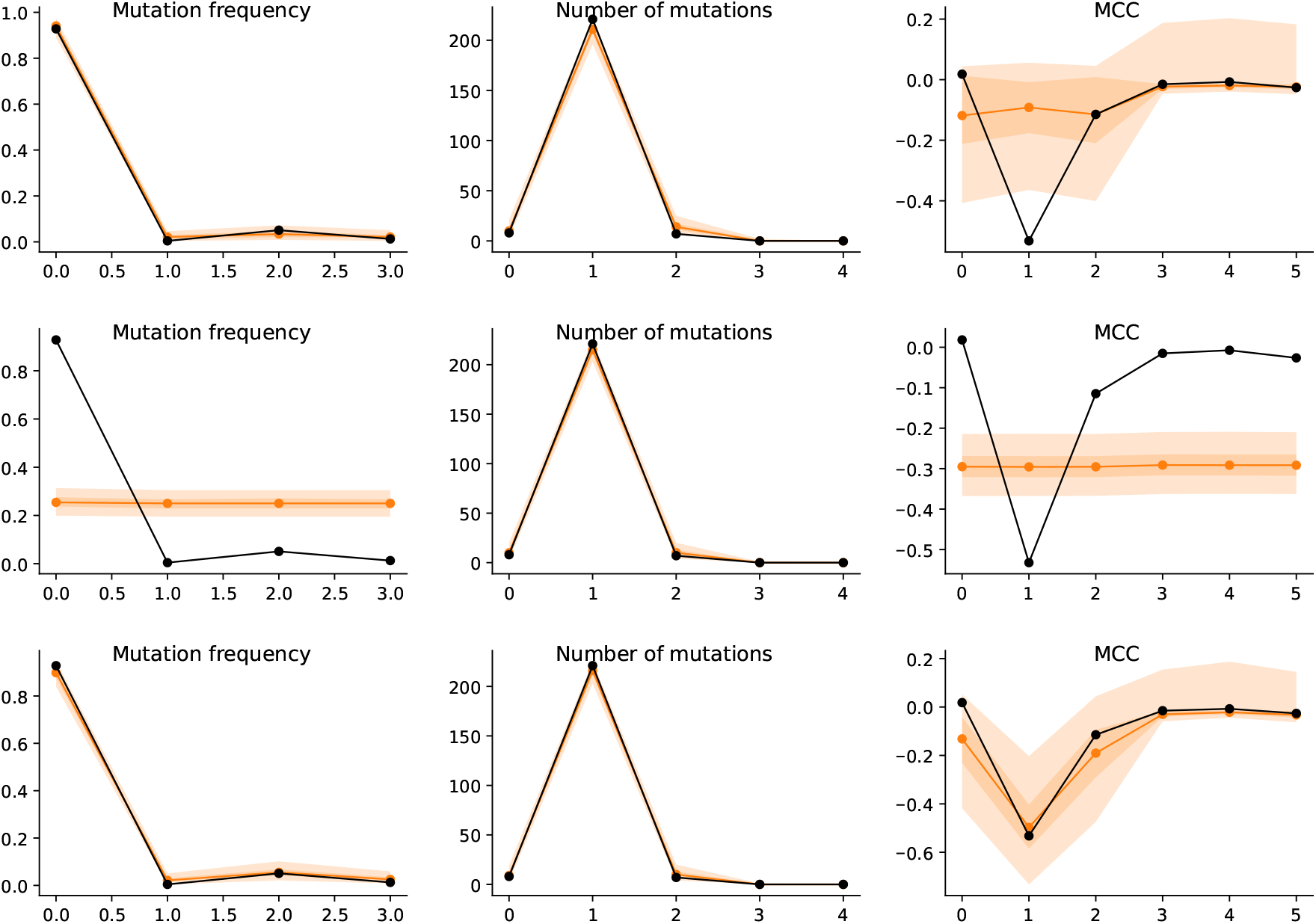
Posterior predictive checks for three models applied to gene set *A*. Top row: the background model. Middle row: rigid mutual exclusivity model (Eq. 2). Bottom row: the flexible mutual exclusivity model (Eq. 3).

The full model (Eq. 4) exhibits convergence issues (what is diagones by 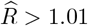) and may have multimodal posterior (Fig. 6). However, accepting that the posterior may not be perfect, this model also suggests that alterations in EGFR and IDH1 may be mutually exclusive.

### B.2 Analysis of the gene set *B*

Fig. 9 shows that the background model underfits positive MCC values. The mutual exclusivity models better capture the underlying distribution. However, as we see in Fig. 8, the mutual exclusivity models do not support mutual exclusivity, rather suggesting possible co-occurrence.

**Figure 8.**
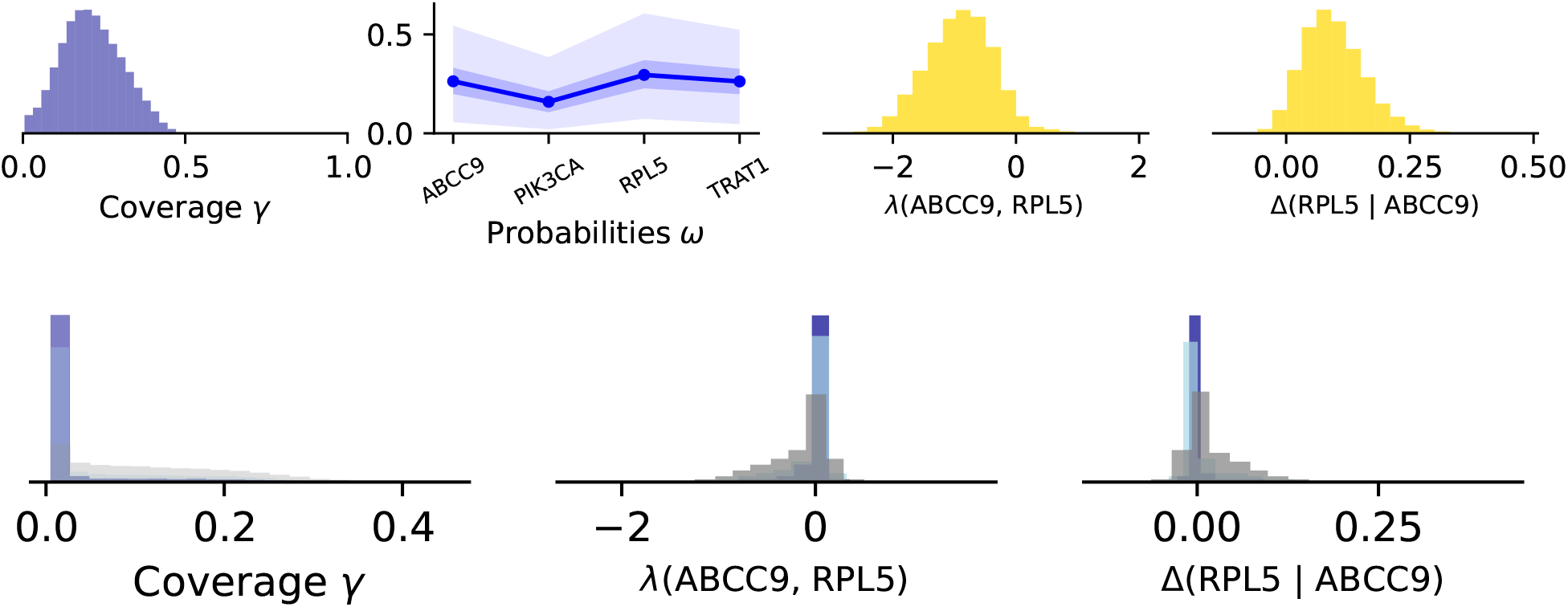
Predictions from different models applied to gene set *B*. Top row: flexible mutual exclusivity model (Eq. 3). Bottom row: full model (Eq. 4) with different priors on the coverage parameter *γ*.

**Figure 9.**
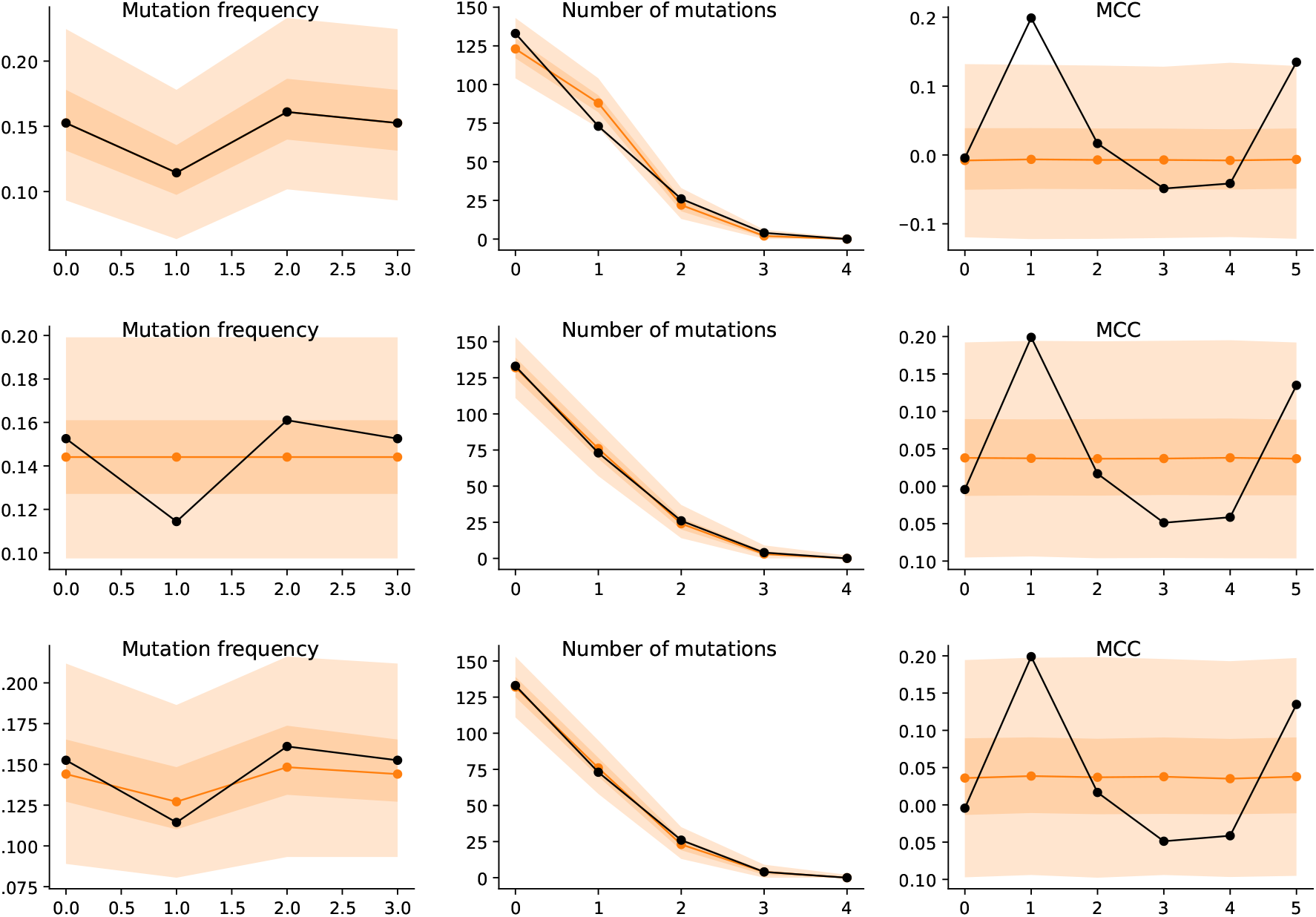
Posterior predictive checks for three models applied to gene set *A*. Top row: the background model. Middle row: rigid mutual exclusivity model (Eq. 2). Bottom row: the flexible mutual exclusivity model (Eq. 3).

**Figure 10.**
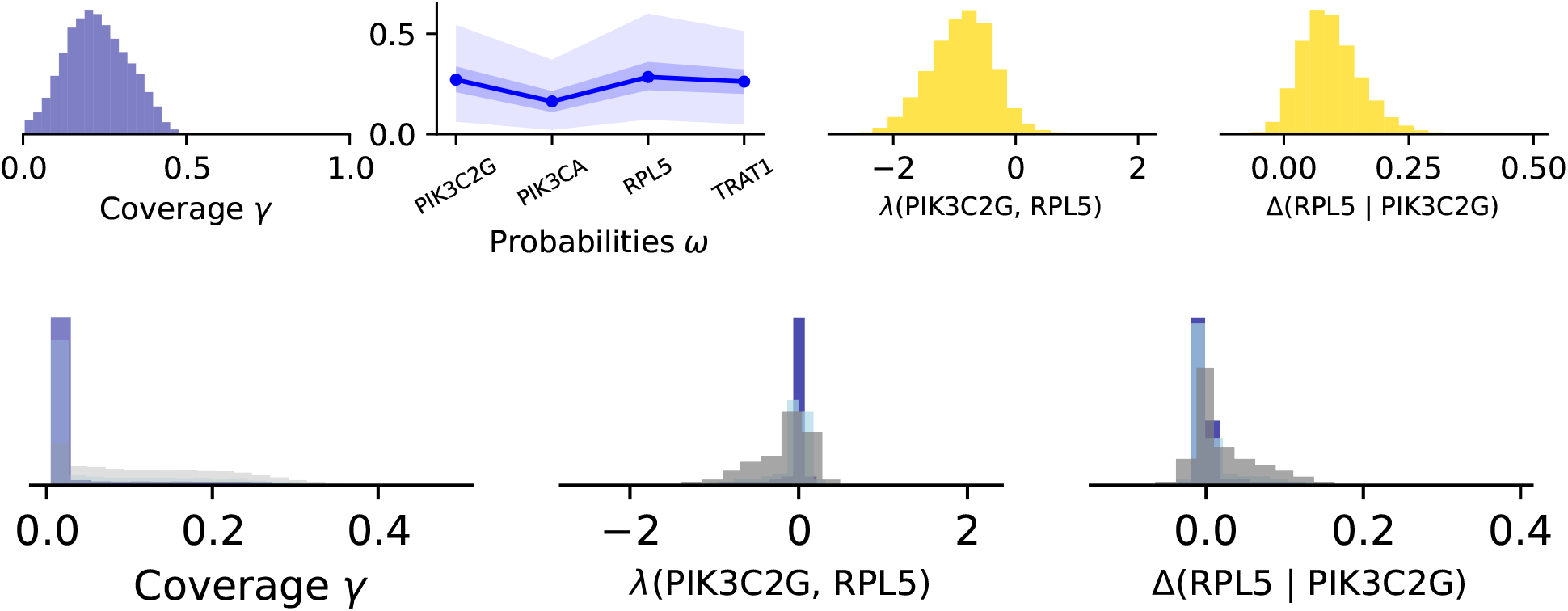
Predictions from different models applied to gene set *B*. Top row: flexible mutual exclusivity model (Eq. 3). Bottom row: full model (Eq. 4) with different priors on the coverage parameter *γ*.

**Figure 11.**
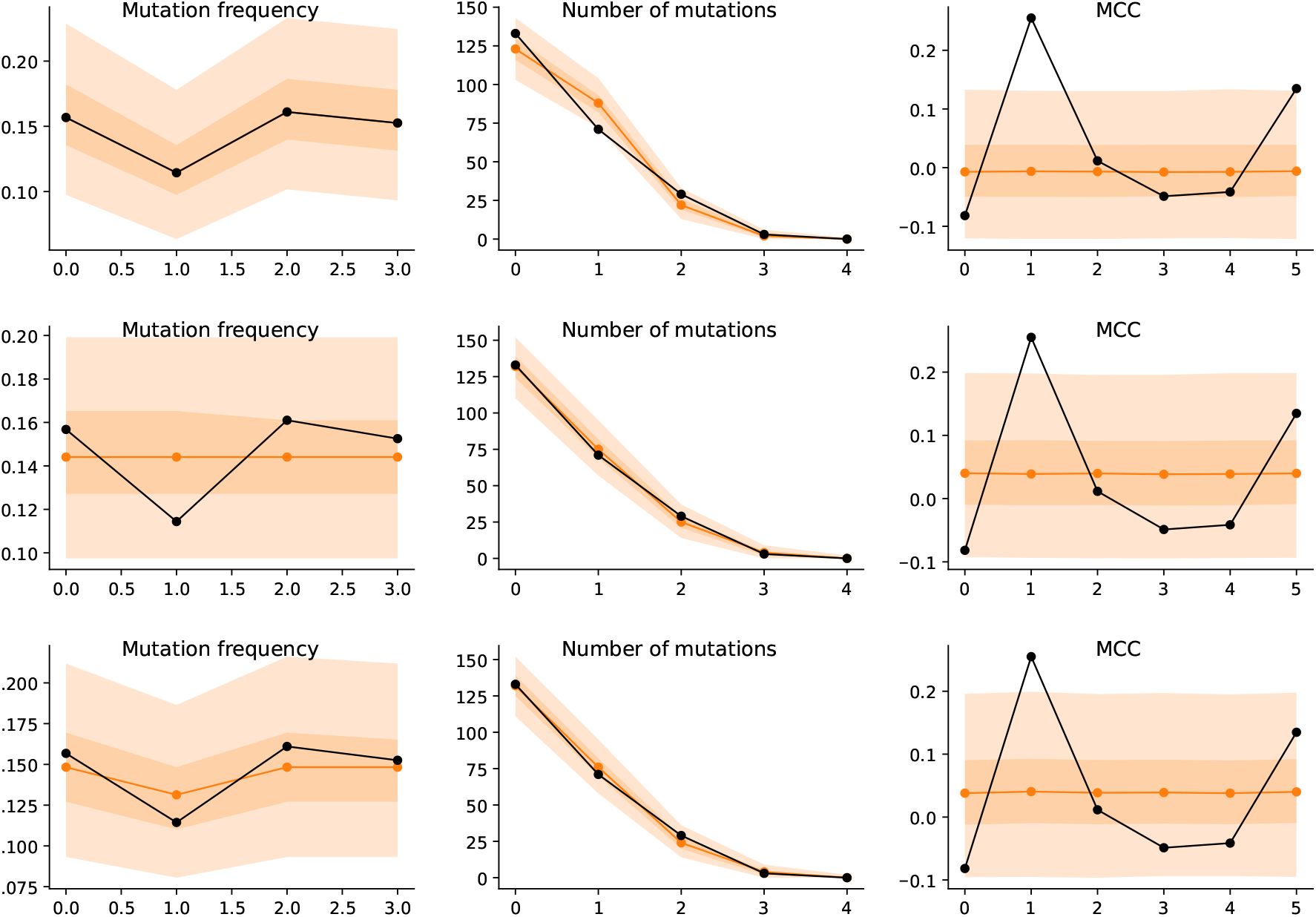
Posterior predictive checks for three models applied to gene set *B*. Top row: the background model. Middle row: rigid mutual exclusivity model (Eq. 2). Bottom row: the flexible mutual exclusivity model (Eq. 3).

**Figure 12.**
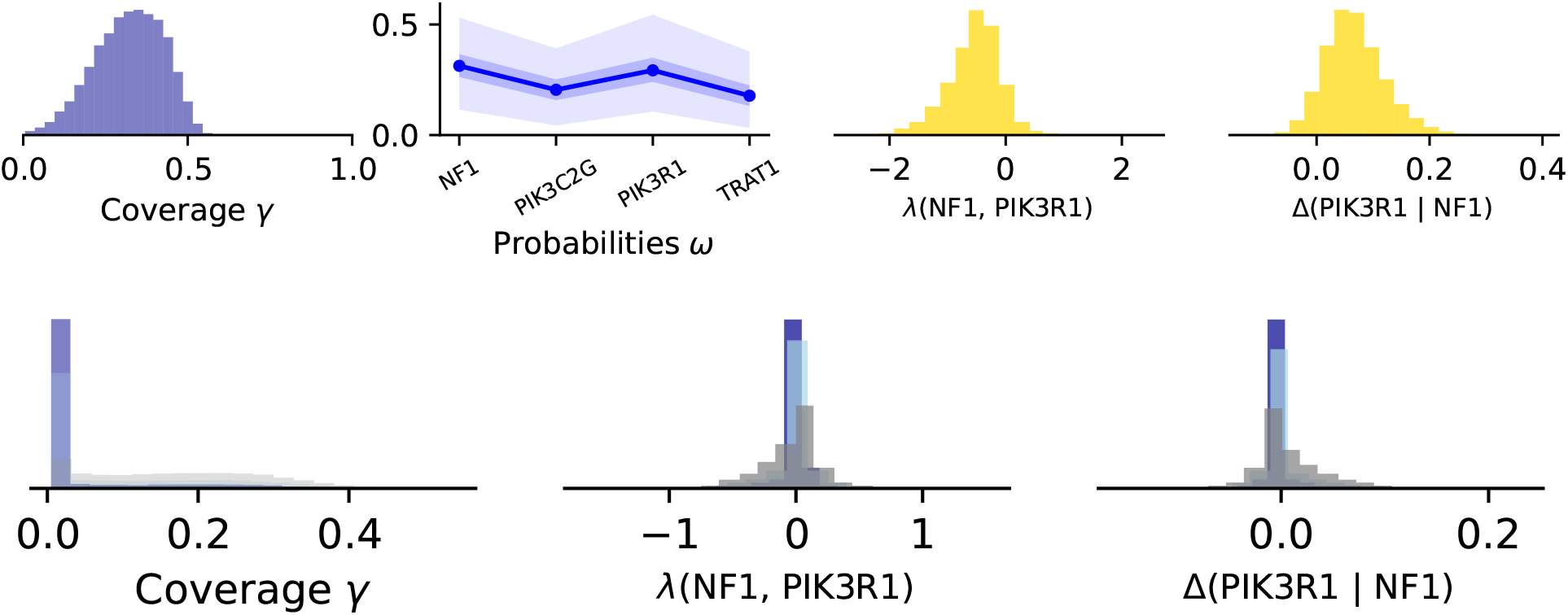
Predictions from different models applied to gene set *C*. Top row: flexible mutual exclusivity model (Eq. 3). Bottom row: full model (Eq. 4) with different priors on the coverage parameter *γ*.

### B.3 Analysis of the gene set *C*

Similarly as in Appendix B.2, the models do not seem to support mutual exclusivity. However, they underestimate positive correlations.

### B.4 Analysis of the gene set *D*

As we see in Fig. 13, backkground model underestimates positive correlation. Mutual exclusivity models seem to describe the data adequately, however they do not support evidence for mutual exclusivity.

**Figure 13.**
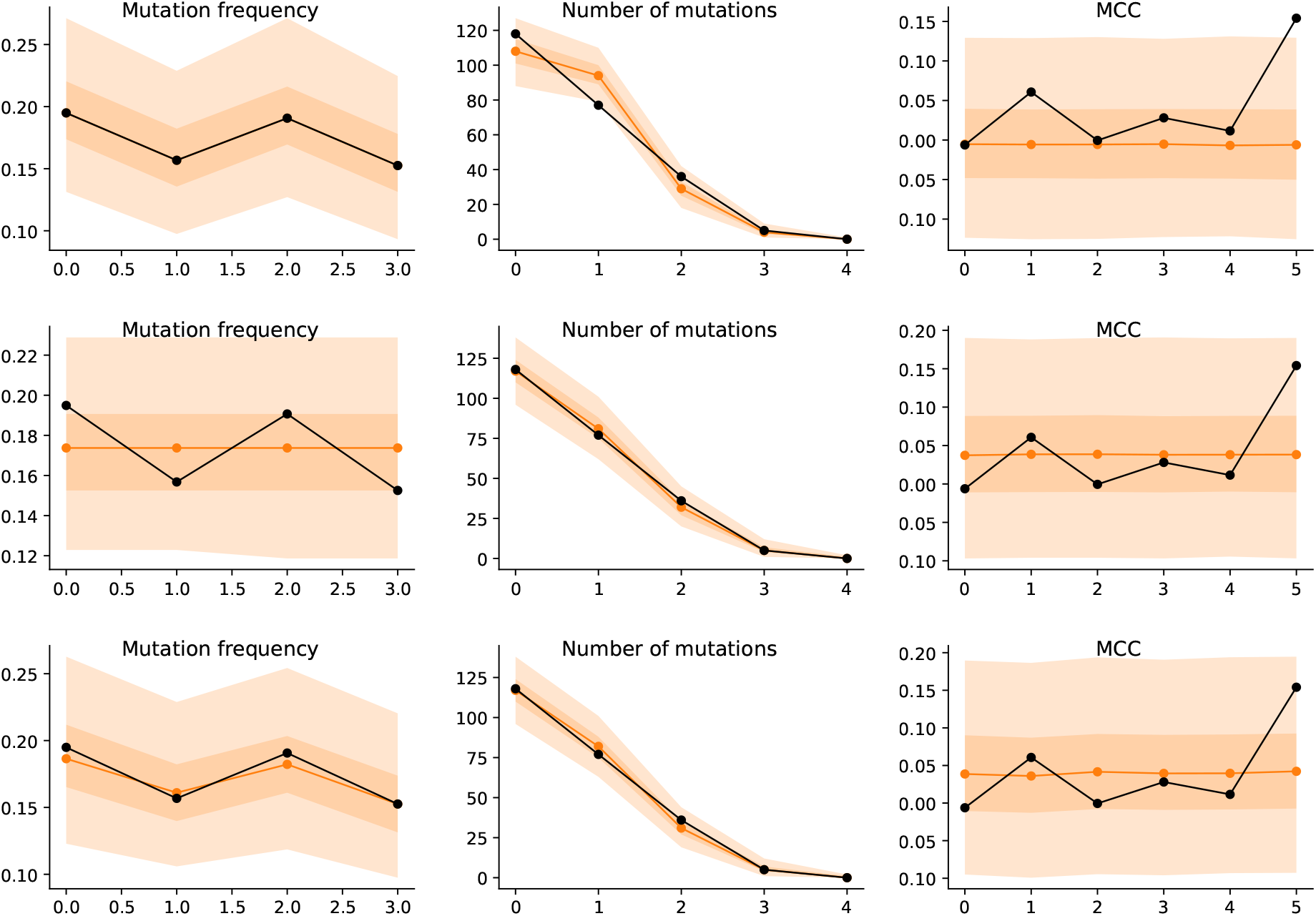
Posterior predictive checks for three models applied to gene set *D*. Top row: the background model. Middle row: rigid mutual exclusivity model (Eq. 2). Bottom row: the flexible mutual exclusivity model (Eq. 3).

## Notes

https://github.com/cbg-ethz/jnotype

### Competing Interest Statement

The authors have declared no competing interest.

